# Cortical and subcortical T1 white/gray contrast, chronological age, and cognitive performance

**DOI:** 10.1101/537043

**Authors:** John D. Lewis, Vladimir S. Fonov, D. Louis Collins, Alan C. Evans, Jussi Tohka, for the Brain Development Cooperative Group, and the Pediatric Imaging, Neurocognition, and Genetics Study

**Affiliations:** Montreal Neurological Institute, McGill University, Montreal, Canada; AIVirtanen Institute for Molecular Sciences, University of Eastern Finland, Kuopio, Finland

**Author notes:** These two authors have contributed equally (John D. Lewis, Jussi Tohka). Corresponding author, Montreal Neurological Institute, NW147, 3801 University Street, Montreal, QC, Canada, H3A2B4.

**Keywords:** brain development, cognitive development, age prediction, cortical white/gray contrast, subcortical white/gray contrast, VIQ-PIQ discrepancies

## Abstract

The maturational schedule of typical brain development is tightly constrained; deviations from it are associated with cognitive atypicalities, and are potentially predictive of developmental disorders. Previously, we have shown that the white/gray contrast at the inner border of the cortex is a good predictor of chronological age, and is sensitive to aspects of brain development that reflect cognitive performance. Here we extend that work to include the contrast at the white/gray border of subcortical structures. We show that cortical and subcortical contrast together yield better age-predictions than any non-kernel-based method based on a single image-type, and that the residuals of the improved predictions provide new insight into unevenness in cognitive performance. We demonstrate the improvement in age predictions in two large datasets: the NIH Pediatric Data, with 831 scans of typically developing individuals between 4 and 22 years of age; and the Pediatric Imaging, Neurocognition, and Genetics data, with 909 scans of individuals in a similar age-range. Assessment of the relation of the residuals of these age predictions to verbal and performance IQ revealed correlations in opposing directions, and a principal component analysis of the residuals of the model that best fit the contrast data produced components related to either performance IQ or verbal IQ. Performance IQ was associated with the first principle component, reflecting increased cortical contrast, broadly, with almost no subcortical presence; verbal IQ was associated with the second principle component, reflecting reduced contrast in the basal ganglia and increased contrast in the bilateral arcuate fasciculi.

## 1. Introduction

Growth charts, *e.g.* those from the Centers for Disease Control and Prevention (https://www.cdc.gov/growthcharts/), are used by pediatricians and parents to track the growth of infants, children, and adolescents. Deviation from the typical growth trajectory is an indicator of potential issues. But such charts provide only measures such as height/length, weight, and head circumference; more detailed measures of brain development can be obtained from brain imaging data. Chronological age can be quite accurately predicted from brain images (Dosenbach et al., 2010; Brown et al., 2012; Franke et al., 2012; Mwangi et al., 2013; Khundrakpam et al., 2015; Ball et al., 2017; Lewis et al., 2018), and deviations from the typical maturational schedule are associated with differences in cognitive functioning (Erus et al., 2015; Lewis et al., 2018). Determining the extent to which typical cognitive development rests upon adherence to the typical maturational trajectory is critical to our understanding of development, and to our ability to utilize brain data to identify atypical development so that we might intervene. Furthermore, to the extent that we can associate atypical cognitive development with specific brain regions, both our understanding and our ability to intervene are enhanced.

In samples representative of normal development from early childhood through adolescence, correlations between chronological age and age estimates from brain images are as high as 0.96 based on multi-metric data from mutiple image types (Brown et al., 2012). Brown et al. (2012) reported substantially lower correlations for a single image-type; for T1-weighted data their multi-metric approach, including cortical and subcortical measures, yielded a correlation between chronological and estimated age of approximately 0.91. Their multi-metric estimates of age from T2-weighted signal intensity also show a correlation with chronological age of 0.91, with contributions from both white-matter tracts and subcortical gray matter (Brown et al., 2012). Estimates of age based on diffusion data show a correlation close to 0.9, with the greatest contributions coming from both cortical and subcortical gray matter, and long-range connections, e.g. the corpus callosum (Mwangi et al., 2013; Erus et al., 2015; Brown et al., 2012). Note that measures from subcortical material contribute to each of these results. Moreover, Brown et al. (2012) showed, albeit in their multi-image-type model, that subcortical volume measures were a core contributor to their predictions. Indeed, the subcortical maturational trajectory differs from that of the cortex, and differs between different subcortical structures (Raznahan et al., 2014). Thus, such measures should boost the accuracy of MRI-based age-prediction models.

Our recent work (Lewis et al., 2018) yielded a correlation between chronological and estimated age of approximately 0.91, using only T1-weighted data, and only a single measure: the intensity contrast between white and gray matter at the inner edge of the cortex. This result is superior to any single-metric method (*e.g.* Khundrakpam et al. (2015) or Dosenbach et al. (2010)), and comparable to the best multi-metric methods using a single image-type (*e.g.* Brown et al. (2012) or Ball et al. (2017)). But it seems likely that this result could be improved upon by including the intensity contrast between white and gray matter at the boundary between subcortical gray matter and the surrounding white matter, which is a novel measure, to the best of our knowledge, but a natural extension of our previous work (Lewis et al., 2018). Moreover, it seems likely that including the intensity contrast at the subcortical gray-white boundaries might capture additional aspects of brain maturation related to cognitive development. Subcortical regions play important roles in cognitive, affective, and social functions (Johnson, 2005; Utter and Basso, 2008; van Schouwenburg et al., 2010; Fischi-Gómez et al., 2014; Humphries and Prescott, 2010; Grazioplene et al., 2015; Bohlken et al., 2016; Guo et al., 2006; Isaacs et al., 2008), and subcortical abnormalities have been associated with various psychiatric disorders (Rimol et al., 2010; Hartberg et al., 2011; Cerliani et al., 2015; Hibar et al., 2016; Hoogman et al., 2017).

Here we test these conjectures. We devise methods to extract surfaces for the caudate, putamen, globus pallidus, and thalamus, and extend our methods for measuring white/gray contrast at the inner edge of cortex to measure white/gray contrast at the gray-white boundaries of these subcortical structures. We then evaluate the impact on age-prediction accuracy of including subcortical white/gray contrast measures in addition to cortical white/gray contrast measures. We do so in two large datasets: the NIH Pediatric Data and the Pediatric Imaging, Neurocognition, and Genetics data. We then assess the relation of the residuals of these age predictions to the verbal and performance IQ measures available in the NIHPD dataset, and map these relationships to the brain.

## 2. Materials and Methods

### 2.1. Data

The data used were taken from two large-scale datasets: the NIH Pediatric data (NIHPD); and the Pediatric Imaging, Neurocognition, and Genetics data (PING). Detailed descriptions of the NIHPD and PING samples are given in (Evans et al., 2006) and (Brown et al., 2012), respectively; here we give a brief summary. Table 1 provides the demographic information for the two populations.

**Table 1:**
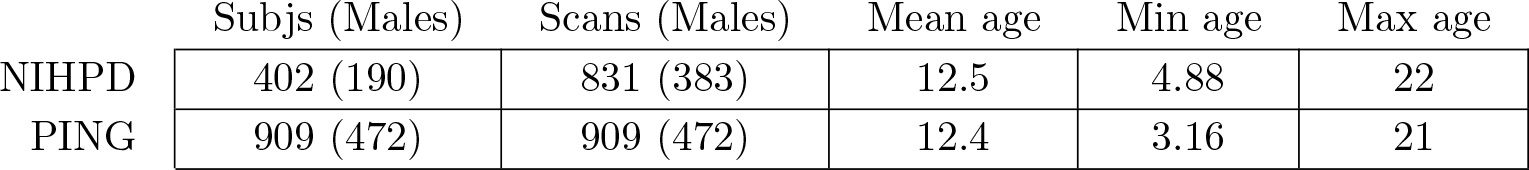
Subject demographics for the two datasets. Ages are given in years.

|  | Subjs (Males) | Scans (Males) | Mean age | Min age | Max age |
| --- | --- | --- | --- | --- | --- |
| NIHPD | 402 (190) | 831 (383) | 12.5 | 4.88 | 22 |
| PING | 909 (472) | 909 (472) | 12.4 | 3.16 | 21 |

#### 2.1.1. The NIHPD sample

The NIHPD data were collected to allow the characterization of healthy brain development, and the relation between brain development and behaviour. Both high-quality MRI data and comprehensive clinical/behavioral measures were collected. Recruitment was epidemiologically based, demographically balanced, and used strict exclusion factors (biasing the sample upward in terms of IQ). The sample includes over 400 children ranging from 4.5 to 18.5 years of age, well-distributed across the range. The database is a mix of longitudinal and cross-sectional data, but we consider all data as cross-sectional, taking care to ensure that the evaluation of predictive models is not biased by this dependence. The data were collected at six sites: Children’s Hospital, Boston; Children’s Hospital Medical Center, Cincinnati; University of Texas Houston Medical School, Houston; Neuropsychiatric Institute and Hospital, UCLA; Children’s Hospital of Philadelphia; and Washington University, St. Louis. Data were acquired on either a General Electric 1.5T scanner or a Siemens Medical Systems 1.5T scanner. Pulse sequences and parameters were chosen to maximize image quality and minimize differences across sites. Each site acquired multiple image-types (T1-weighted, T2-weighted and PD-weighted); the current study utilizes only the T1-weighted data, which was acquired with a 3D T1-weighted spoiled gradient recalled echo sequence with a resolution of 1mm isotropic.

#### 2.1.2. The PING sample

The PING dataset includes data from more than 900 typically developing children between the ages of 3 and 20 years, including individuals with learning or language disabilities. Subjects were excluded if they had a history of major developmental, psychiatric, and/or neurological disorders, or medical conditions that affect neurological development, or had had a brain injury. As opposed to the NIHPD sample, these data are strictly cross-sectional. Data were collected at 10 sites: Weil Cornell Medical College, University of California at Davis, University of Hawaii, Kennedy Krieger Institute, Massachusetts General Hospital, University of California at Los Angeles, University of California at San Diego, University of Massachusetts Medical School, University of Southern California, and Yale University. Data were acquired on either a General Electric 3T scanner, a Siemens Medical Systems 3T scanner, or a Philips 3T scanner. Pulse sequence parameters were chosen to maximize image quality and minimize differences across scanners. Across sites and scanners, a standardized multiple image-type high-resolution structural MRI protocol was implemented. Each site acquired T1-, T2-, and diffusion-weighted scans; the current study uses only the T1-weighted data, acquired with a 3D T1-weighted inversion prepared RF-spoiled gradient echo sequence with a resolution of 1mm isotropic, using prospective motion correction. Specific protocols for each scanner manufacturer are provided at ^1^

### 2.2. Surface measurements

#### 2.2.1. Cortical surface extraction

The T1-weighted volumes were processed with CIVET (version 2.1; 2016), a fully automated structural image analysis pipeline developed at the Montreal Neurological Institute ^2^. CIVET corrects intensity non-uniformities using N3 (Sled et al., 1998); aligns the input volumes to the Talairach-like ICBM-152-nl template (Collins et al., 1994); classifies the image into white matter, gray matter, cerebrospinal fluid, and background (Zijdenbos et al., 2002; Tohka et al., 2004); extracts the white-matter and pial surfaces (Kim et al., 2005); and maps these to a common surface template (Lyttelton et al., 2007).

#### 2.2.2. Subcortical surface extraction

Subcortical segmentation into left and right caudate, putamen, globus pallidus, and thalamus was done using a label-fusion-based labeling technique based on Coupé et al. (2011) and further developed by Weier et al. (2014), incorporating ideas from Landman et al. (2012). The approach used a population-specific template library. In the current work, the library for each dataset was constructed by clustering (as described below) the deformation fields from the non-linear transforms produced by CIVET, and using the central-most subject of each cluster to construct the entries in the template library. Thus, the template library represented the range of deformations found in the population.

As the basis for clustering, the Jacobian was computed for the non-linear transform produced by CIVET for each subject. The values in the Jacobian were then extracted as a vector for each voxel within a mask formed by binarizing the template-based subcortical sementation, eroding it 1 mm and dilating it 5 mm in order to capture the anatomical context of the nonlinear registration in this area. These Jacobian vectors were then clustered using an equal combination of cosine similarity and Euclidean distance with Ward’s clustering method (Ward Jr, 1963), with the number of clusters chosen as the square of the natural log of the number of subjects. Then, within each cluster, the sum-squared distance from each subject to each other subject was computed, and the subject with the minimum sum-squared distance was taken as the central-most subject of the cluster.

To create the library entry for a cluster, the non-linear transform for the central-most subject was inverted and used to warp the ICBM-152-nl template together with the subcortical segmentation defined on it; this pair was then added to the template library. The template library was thus a set of warped copies of the ICBM-152-nl template together with their correspondingly warped segmentations. Once the template library had been created, each subject in the population was non-linearly registered to the *n* closest templates in the library (here, *n* = 7), and the resulting transforms were used to warp their corresponding segmentations to the subject; the final labelling was then established via a per-voxel majority vote.

Once the subcortical structures for a subject were labeled, surfaces were fit to these labels. The surfaces for each structure were created on the ICBM-152-nl template. These model surfaces were warped to each individual based on the transforms derived from the label-fusion-based labeling stage, and adjusted to the final labels by moving vertices along a distance map created for each label. The surfaces for each structure were then registered to their corresponding common surface template to ensure cross-subject vertex correspondence, as per the cortical surfaces.

#### 2.2.3. White/gray contrast measurements

To extract the white/gray contrast measures, the intensity on the T1-weighted MRI was sampled inside and outside of the surface at the (cortical or subcortical) gray-white boundary, and the ratio of the two measures was formed. More specifically, for the cortex, the gray matter was sampled 1 mm outside of the white surface, and white matter was sampled 1 mm inside of the white surface. These choices accommodate both the thinnest areas of the cortex and very thin gyri. For the subcortical structures, due to the absence of issues related to the thickness of either the gray matter or the surrounding white matter, and to the lesser clarity of the boundary between white and gray, the white matter was sampled 2 mm outside of the subcortical white surface, and gray matter was sampled 2 mm inside of the subcortical white surface. An example of the surfaces used to form the white/gray contrast measures is provided in Figure 1.

In more detail, a distance map was created from the white surface, smoothed with a Gaussian kernel, and used to create a gradient vector field. At low resolution, issues arise with this vector field in areas with very thin strands of white matter, e.g. at the tips of gyri, and so the distance maps were created at a resolution of 0.25 mm and smoothed with a 0.5 mm FWHM Gaussian kernel before creating the gradient vector field. The cortical white surface was moved 1mm inward along this gradient vector field to produce a sub-white surface, and 1 mm outward to produce a supra-white surface. The intensity values on the T1-weighted image (without non-uniformity correction or normalization) were sampled at each vertex of both the supra-white surface and the sub-white surface, and the ratio was formed by dividing the value at each vertex of the sub-white surface by the value at the corresponding vertex of the supra-white surface. The same procedure produced the contrast measures for the subcortical surfaces, but moving inward and outward 2 mm instead of 1 mm.

**Figure 1:**
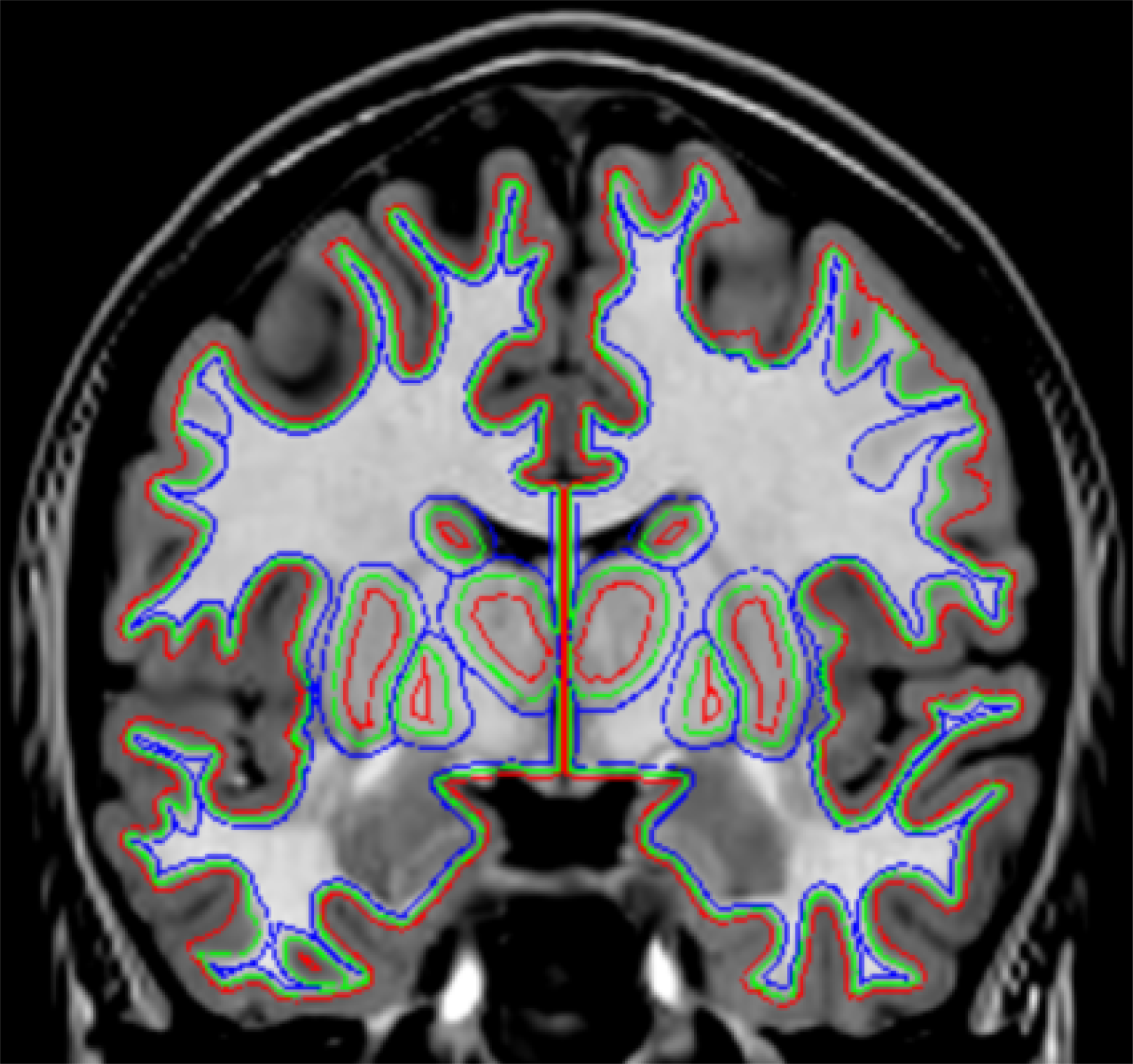
An example of the surfaces used to form the the white/gray contrast measures. The surface at the gray-white border is shown in green. This surface is extracted by CIVET, for the cortex, and by fitting surfaces to the labels created by label-fusion-based methods, for the subcortical labels. A distance map and corresponding vector field were created from this, and copies of the surfaces were moved into gray matter (red) and white matter (blue) along the gradient vectors of this distance map. For the cortex, both deriviative surfaces were moved 1mm along this vector field; for the subcortical structures, both derivative surfaces were moved 2mm along this same vector field. Gray matter and white matter intensity were then measured at each of the vertices of these derivative surfaces, and the white/gray contrast measure formed as the ratio of white intensity to gray intensity at corresponding vertices of the two surfaces.

Since the intensities that form the ratio are sampled very near to one another, the effects of image inhomogeneities will be small. But, the white/gray contrast measures are sensitive to scanner-specific differences in tissue contrast. To correct for this, we normalized the contrast values per scanner manufacturer. The normalized contrast measure is 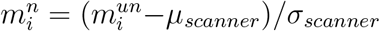, where 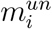 is the unnormalized contrast measure and *μ*_*scanner*_ and *σ*_*scanner*_ are the mean and the standard deviation of the contrast values across the surface and across all of the subjects whose data were acquired on *scanner*. This correction is equivalent to scanner-wise z-scoring. This correction was done separately for each of the surfaces (the two cortical surfaces and the eight subcortical surfaces). Left and right surfaces were considered together. The effectiveness of this correction was demonstrated in (Lewis et al., 2018). For the rest of the paper, we drop the superscripts and always refer to the normalized contrast.

### 2.3. Age prediction

Our age prediction method and its evaluation are unchanged from (Lewis et al., 2018), but we summarize the method to make this paper more self-contained. The original 81,924 measurements on the cortical surface were grouped into 640 parcels by recursively merging the neighbouring triangles of the surface mesh model, yielding regions of approximately equal surface area. We have previously studied predicting age with different number of parcels and concluded that 640 parcels yielded the most accurate age predictions (Lewis et al., 2018). The thalamus, caudate, putamen, and globus pallidus were generated with far fewer vertices, and so we grouped their measurements into 160 parcels per structure in the same way as for the cortical surface in order to attain parcels with approximately the same surface area as the cortical parcels. This resulted in 640 parcels also for the subcortical contrast measures. We assumed a linear model for predicting subjects’ ages based on the white/gray contrast measurements. The model is

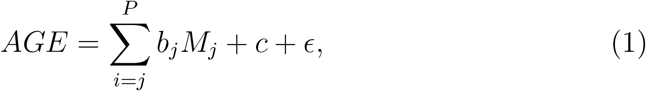

where AGE is the age of the subject (in days); *m*_*j*_, *j* = 1 … *P*, are the contrast measurements for each parcel; *b*_*j*_ and *c* are the model weights to be learned by the machine learning algorithm, and *∈* is an error term. We considered age prediction with subcortical contrast alone, with cortical contrast alone, and with both together. Subcortical contrast was formed by concatenating the contrast measures from the eight subcortical surfaces to form a vector of length 640. We estimated the parameters **b** = [*b*_1_, …, *b*_*P*_] and *c* of the model by penalized least squares with the elastic-net penalty as implemented in the Glmnet package (Friedman et al., 2010) ^3^. This equals to the minimization of the cost function

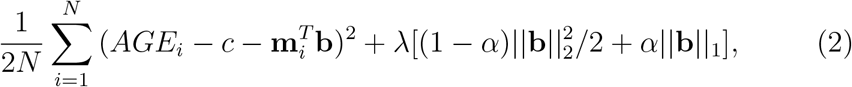

where subscript *i* refers to scans, **m**_*i*_ = [*m*_*i*1_, …, *m*_*iP*_] are the measurements for scan *i* and *N* is the number of scans. The elastic-net penalty 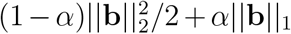 is a weighted average of the LASSO and ridge penalties (Zou and Hastie, 2005). The LASSO penalty ‖**b**‖_1_ forces many parameters to have a zero-value leading to the variable selection while the ridge penalty 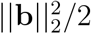 helps to ensure that highly correlated variables are selected simultaneously and they have similar model weights (Zou and Hastie, 2005). We set *α* = 0.5, as in Khundrakpam et al. (2015) and Lewis et al. (2018). The relative weight of the data term and the penalties, denoted by the parameter *λ* ∈ ℝ in Eq. (2), was decided based on cross-validation from a sequence of 300 values decreasing on the log scale (Friedman et al., 2010). We standardized the input measurements so that 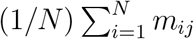 and 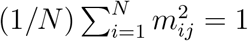 for *j* = 1, …, *P*. Thus, the parameter estimates *b*_*j*_ were also standardized (called standardized regression coefficients) and more comparable to each other than the unstandardized coefficients, see *e.g.* Gelman (2008). The logic of standardized coefficients is to re-express *b*_*j*_ as the effect of a one-SD change in *M*_*j*_ as opposed to a unit change in *M*_*j*_.

### 2.4. Evaluation of age prediction

We selected the *λ* and evaluated the age predictions using two nested stratified 10-fold cross-validation (CV) loops (Ambroise and McLachlan, 2002; Huttunen et al., 2012) ^4^. The value for the parameter *λ* was selected in the inner CV loop by minimizing the mean squared error among the candidate values, and the age predictions were evaluated in the outer CV loop, thus avoiding the training on testing data problem. We repeated the nested CV loops 10 times to reduce the random variation in the evaluation of accuracy due to the random selection of the folding scheme. The model goodness criteria applied were the correlation coefficient between the chronological and estimated age, and the mean absolute error (MAE) between the chronological and estimated age. For the correlation coefficient, we averaged the 10 distinct correlation values stemming from the 10 CV runs. Since with the NIHPD sample we had up to three sets of measurements of certain subjects at different ages, we controlled for the non-independence of the observations in the CV by placing all the scans of the subject *i* in the same fold and thus all scans of a given subject were either in the training set or in the test set. Therefore, data from subject *i* was never used to build an age-predictor for subject *i*. This is an important consideration and failure to account for non-independence would lead to positively biased model accuracy estimates.

As described in (Lewis et al., 2018), we computed 95% confidence intervals (CIs) for cross-validated correlations using a bootstrap method and compared the cross-validated MAEs from the different sets of measurements using a permutation test.

### 2.5. Evaluation of relation of residuals to cognitive functioning

With the NIHPD database there are IQ scores for a subsample of 760 scans from 391 subjects. Previously we showed that the residuals of our age predictions based on cortical contrast, defined as *chronological age – predicted age*, were related to cognitive functioning (Lewis et al., 2018). Here, we looked at these relationships in more detail. First, we assessed the relationship of the age-prediction residuals to the two aspects of full scale IQ, *i.e.* performance IQ and verbal IQ, in each case assessing one while controlling for the effect of the other on age prediction, as well as the effect of brain volume, within a linear mixed-effects model. Then, we decomposed the residuals of the model that best fit the contrast data, and assessed the relation of the top two components to performance IQ and verbal IQ.

The best-fit model was determined by searching all possible models comprised of terms including any of ‘AGE’, ‘SEX’, ‘BRAINVOL’, ‘SITE’, ‘AGE*AGE’, and ‘AGE*SEX’, as well as the terms required for a mixed effects model, *i.e.* ‘random(subjects) + I + 1’, where ‘random(subjects)’ induces equal correlations between observations on the same subject, and ‘I’ allows for independent “white” noise in every observation. Each model was assessed using the SurfStat toolbox (http://www.math.mcgill.ca/keith/surfstat/) and the Akaike information criterion (AIC) (Akaike, 1976). The best-fit model was defined as the one with the lowest AIC. The best-fit model was then evaluated with *SurfStatLinMod* to produce the coefficients which were then passed to *SurfStatPCA* in order to obtain the first two principal components of the residuals of the model. The correlation between both of these two principal components and both of performance and verbal IQ was then assessed to determine if these measures of cognitive functioning were prominent sources of variance in the residuals of the best-fit model.

## 3. Results

### 3.1. Age prediction accuracy

The cross-validated accuracies of the age predictions are reported in Table 2 and plotted in Figure 2 with significant differences in mean absolute errors (MAE) indicated. For either dataset, the inclusion of subcortical contrast in addition to cortical contrast produced significantly more accurate predictions than either alone. For the 1.5T NIHPD data, the age predictions based on cortical contrast were significantly more accurate than those based on subcortical contrast. For the 3T PING data, subcortical contrast produced predictions that were not significantly different from those produced by cortical contrast. These results are visible in the scatter plots of predicted age versus chronological age (Figure 3). The plots for the predictions based on subcortical contrast show greater variance than do those based on cortical contrast; and those based on the combination of measures show even less variance. And notice that for both datasets, and for subcortical contrast, cortical contrast, and the combination of measures, the age estimates are biased by neither scanner manufacturer nor gender.

**Table 2:** The cross-validated accuracies of age predictions based on subcortical and cortical white/gray contrast, and both measures together. The average cross-validated correlation value (*R*) for each is given, as well as the mean absolute error (*MAE*) in days.

|  | <u>Subcortical</u> |  | <u>Cortical</u> |  | <u>Subcortical + Cortical</u> |  |
| --- | --- | --- | --- | --- | --- | --- |
|  | *R* | *MAE* | *R* | *MAE* | *R* | *MAE* |
| NIHPD | 0.81 | 660 | 0.88 | 542 | 0.89 | 500 |
| PING | 0.90 | 632 | 0.91 | 613 | 0.93 | 540 |

**Figure 2:**
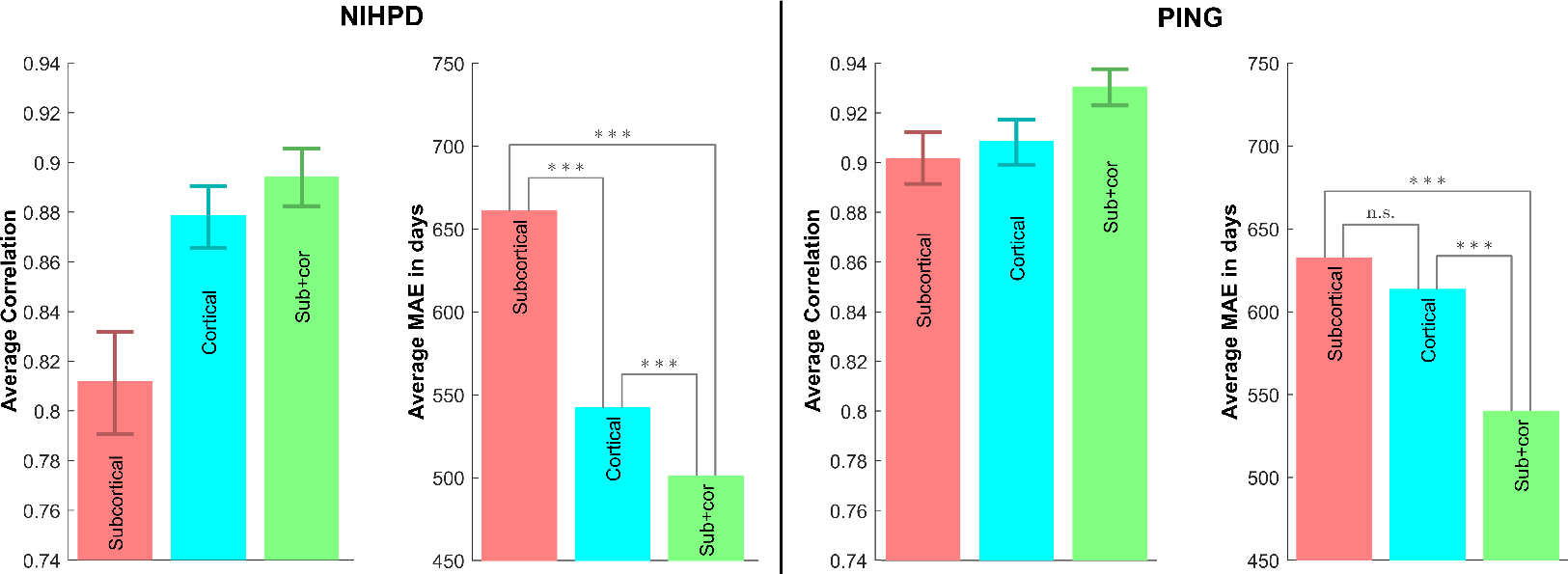
The cross-validated accuracies of the age predictions based on subcortical and cortical white/gray contrast, and both measures together. Glmnet was used as the machine learning algorithm. The correlation plots show, in addition to the average cross-validated correlation value, the 95 % confidence intervals obtained by a bootstrap method. The MAE plots show, in addition to the average cross-validated MAE, a permutation test based comparison between the different methods: n.s stands for not significant, * for *p* < 0.05 ** for *p* < 0.01 and *** for *p* < 0.001.

### 3.2. Age predictors

Our usage of an elastic net penalized linear regression model (Friedman et al., 2010) reveals the brain regions which contribute most to age-prediction. The cortical parcels for which the combined cortical and subcortical white/gray contrast measures reliably contribute to age prediction are shown in Figure 4, for both databases. We defined the signed importance as the median value of weight of the parcel *i* (*b*_*i*_) in the linear regression model across the 10 × 10 cross-validation; thus a parcel had to be selected in the age model at least 50 times during the 10 × 10 cross-validation runs to achieve a non-zero value. The figure shows the non-zero signed importances of all parcels for both databases. Across both databases the non-zero signed importances are broadly distributed across the cortex and the subcortical structures; they are found in each lobe and in each subcortical structure. Thus the age predictions were constructed from information throughout the cortex and the subcortical structures.

**Figure 3:**
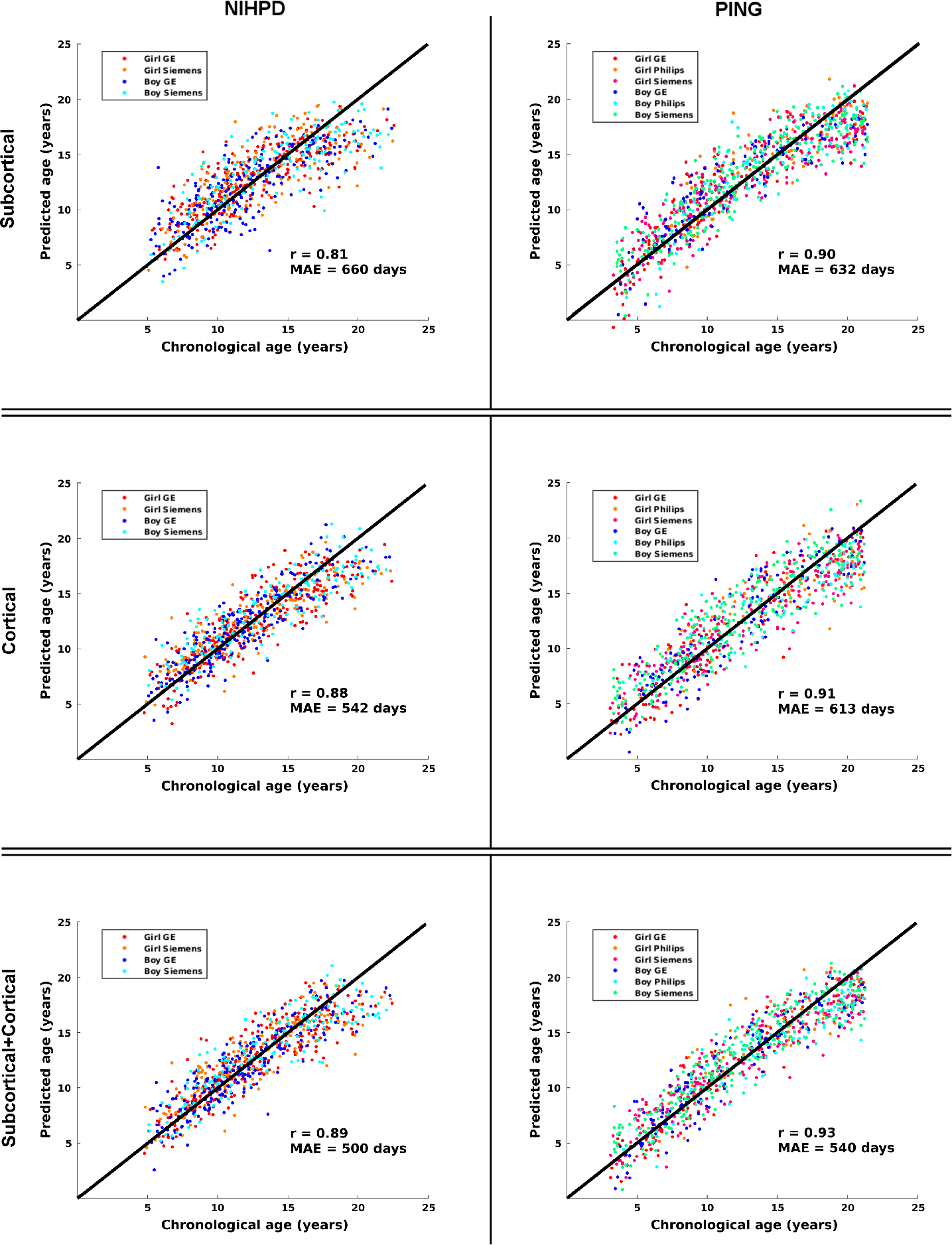
Scatter-plots of predicted versus chronological age during a single cross-validation run (one out of ten) for subcortical contrast alone, for cortical contrast alone, and for subcortical and cortical contrast combined. The black line has a slope of 1 and originates at the origin; this line depicts the optimal predictions. The scanner manufacturer/gender combinations are shown with different colors. Note that there is no apparent bias either due to gender or scanner manufacturer, and that prediction error is relatively uniform across the age span. Note also the lesser spread of the predictions with subcortical and cortical contrast combined than with either alone.

**Figure 4:**
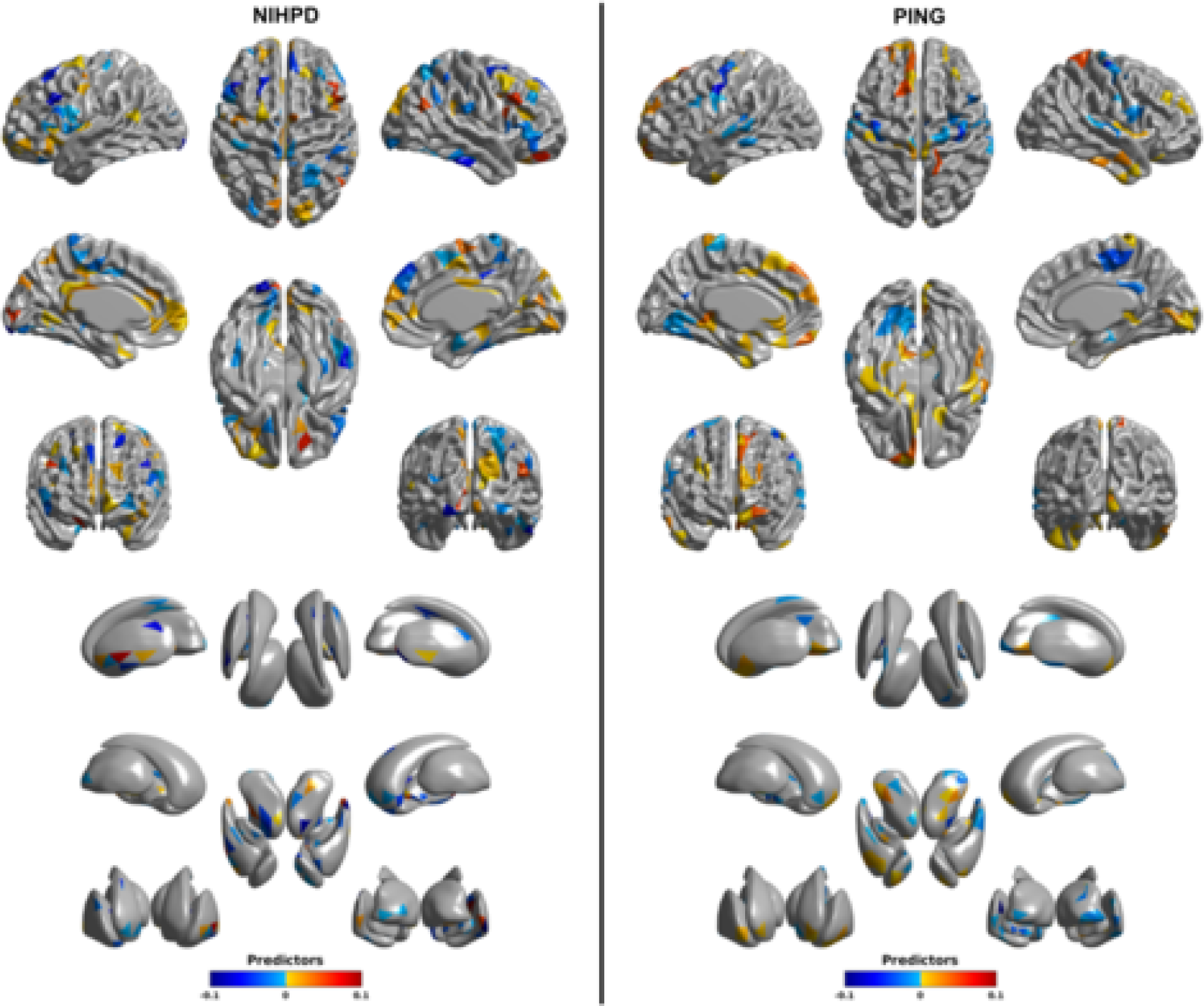
The signed importance of different parcels for age predictions for combined cortical and subcortical white/gray contrast in NIHPD and PING. The signed importance is defined as the median value of weight *b*_*i*_ across the 10 × 10 cross-validation; thus a parcel has to be selected in the age model at least 50 times during the 10 × 10 cross-validation runs to show a non-zero value in the plots. The weights were computed using standardized data so their values are comparable across cortical and subcortical parcels.

### 3.3. Residuals and IQ

The mixed-effects linear model assessing the relationship between the age-prediction residuals and performance IQ (PIQ), controlling for verbal IQ (VIQ) and brain volume revealed a significant positive relationship (*p* < 0.00002). The mixed-effects linear model assessing the relationship between the age prediction residuals and VIQ, controlling for PIQ and brain volume revealed a significant negative relationship (*p* < 0.0174). These relationships are shown in Figure 5.

To map these relationships with IQ to the brain we determined the best-fit mixed-effects linear model explaining white/gray contrast, then decomposed the residuals of that model, and assessed the relation of the top two components to performance IQ and verbal IQ. The model with the lowest AIC among all the possible models comprised of any of ‘AGE’, ‘SEX’, ‘BRAINVOL’, ‘SITE’, ‘AGE*AGE’, and ‘AGE*SEX’, together with ‘random(subjects) + I + 1’ was:

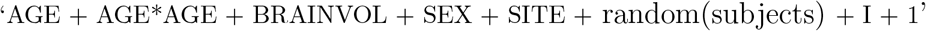

*SurfStatLinMod* together with *SurfStatPCA* were used to obtain the first two principal components of the residuals of this model. The first principal component, explaining 13.0% of the variance, is shown in Figure 6 mapped to the cortical and subcortical surfaces; the second principal component, explaining 3.8% of the variance, is shown in Figure 7. In addition to the first and second principal components, the figures also show the relation of these components to each of performance IQ and verbal IQ. The first principal component involves most of cortex, with the exception of the bilateral pre- and post-central gyri, the calcarine fissure, and the perihippocampal gyri. The subcortical structures are almost completely uninvolved. This component is related to performance IQ, but not verbal IQ. The second principal component involves the cortex adjacent to the bilateral arcuate fasciculi, the middle temporal gyri, and in the opposite direction, the basal ganglia, but not the thalamus. This component is related to verbal IQ, but not performance IQ.

**Figure 5:**
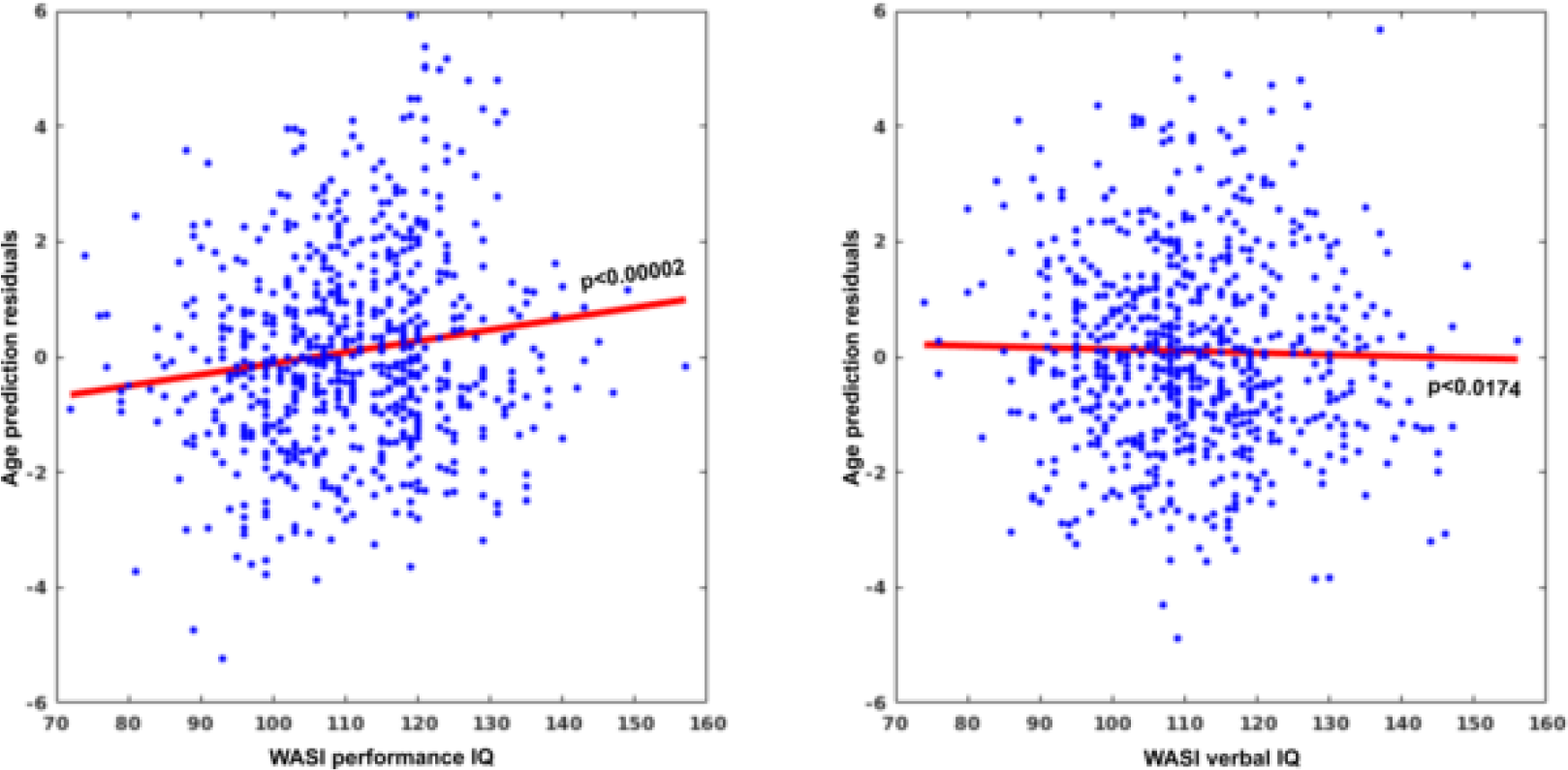
The relation between the age prediction residuals and both PIQ and VIQ. The left scatterplot shows the relation between the age prediction residuals and PIQ, controlling for VIQ and brain volume within a mixed affects model; it is significant positive relationship (p<0.00002). The right scatterplot shows the relation between the age prediction residuals and VIQ, controlling for PIQ and brain volume within a mixed effects model; it is a significant negative relationship (p<0.0174).

**Figure 6:**
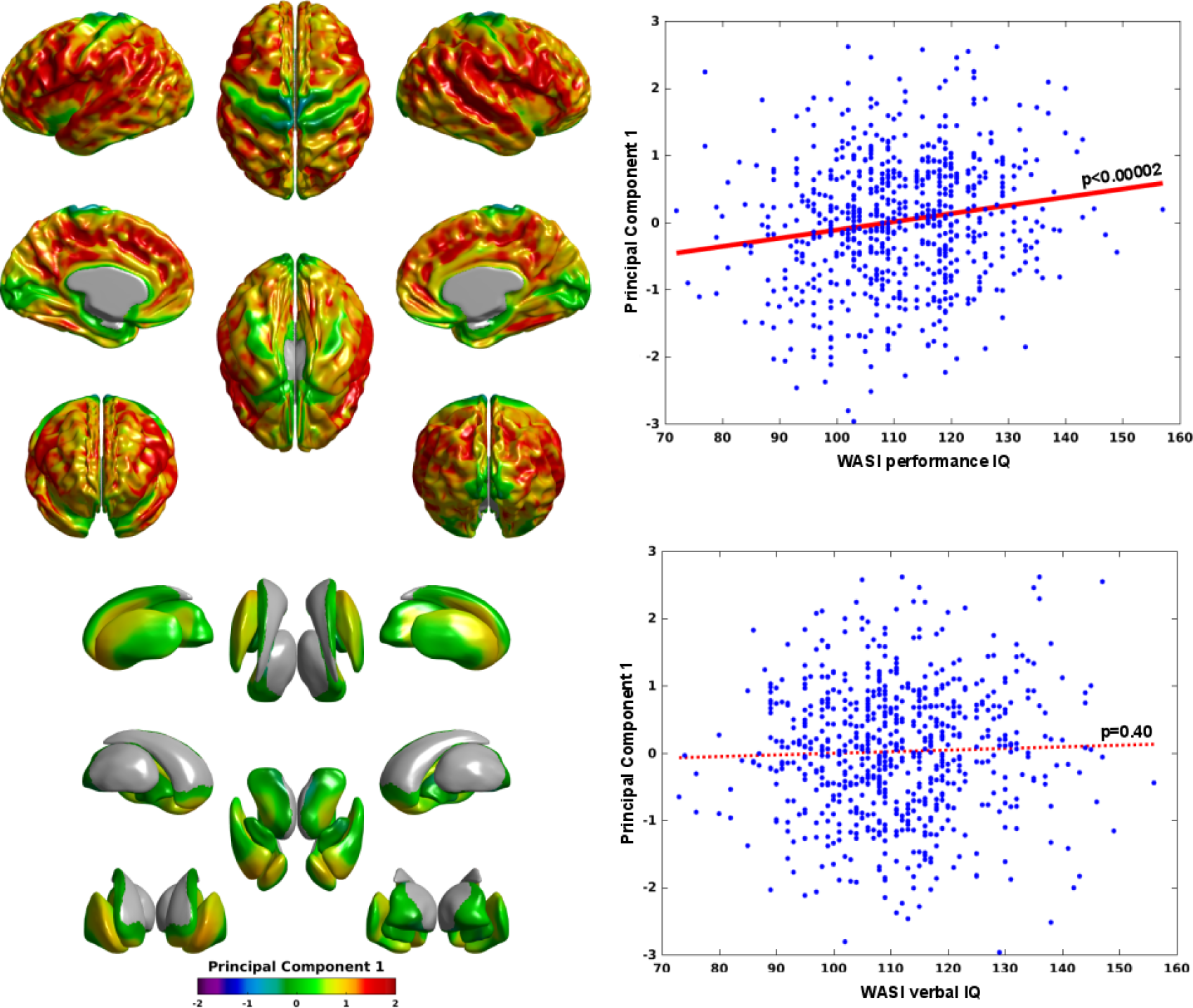
The first principal component and its relation to performance IQ and verbal IQ. Regions in which white/gray contrast measures cannot be generated (due to *e.g.* adjacency to the ventricles) are shown in gray. This component captures variation in cortical contrast, broadly, but not in non-primary areas nor the basal ganglia nuclei of the cerebrum, nor the thalamus. This component is significantly related to performance IQ (p<0.00002) but not verbal IQ.

**Figure 7:**
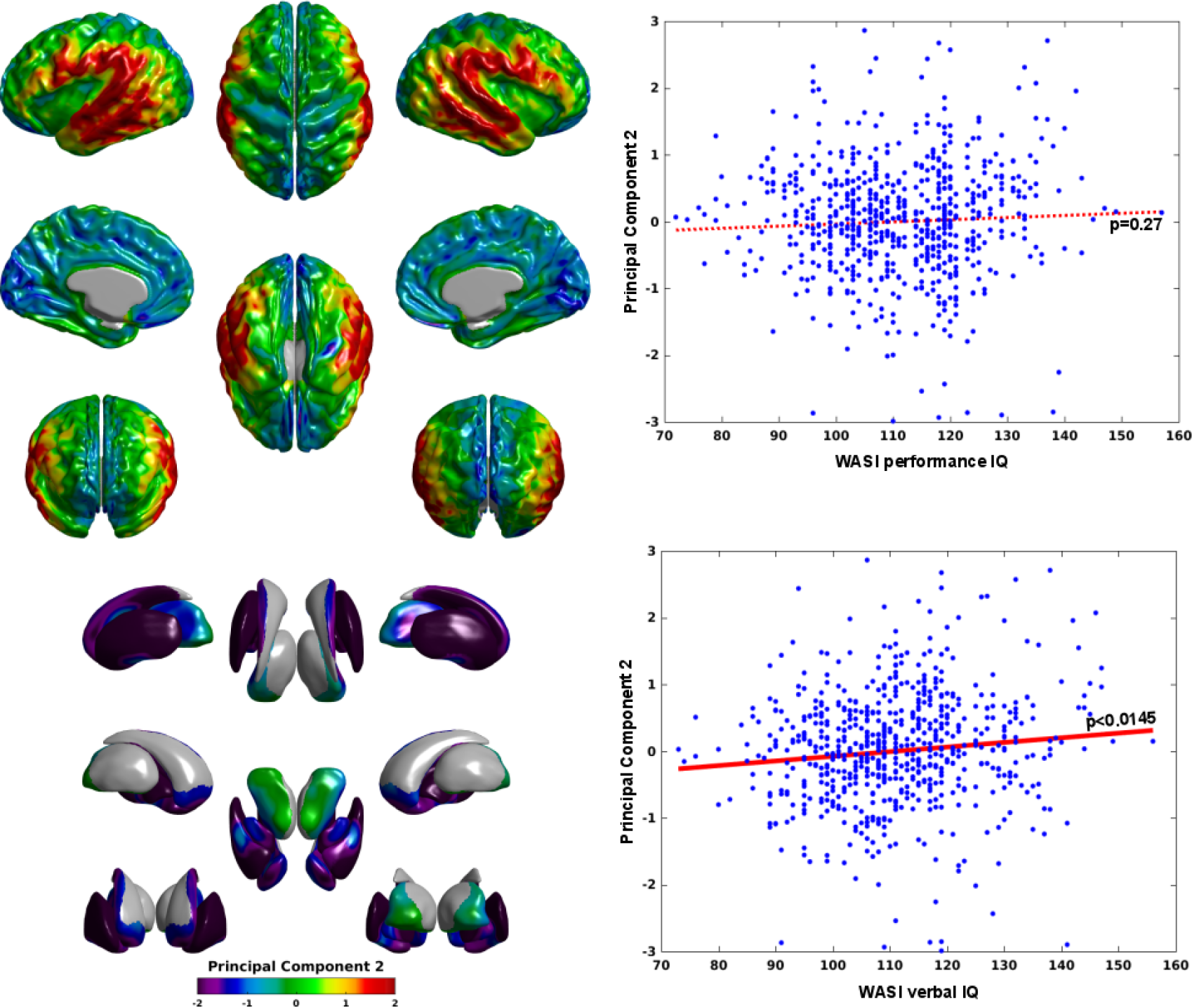
The (inverted) second principal component and its relation to performance IQ and verbal IQ. Regions in which white/gray contrast measures cannot be generated (due to *e.g.* adjacency to the ventricles) are shown in gray. This component involves the basal ganglia nuclei of the cerebrum (but not the thalamus) and inversely the bilateral arcuate fasciculi. It is significantly related to verbal IQ (p<0.0145) but not performance IQ. Note that this is the inverted version of the second principal component; the actual relation of the second principal component to verbal IQ is negative. The inversion makes the interpretation more natural: higher verbal IQ corresponds to greater contrast in the bilateral arcuate fasciculi and lesser contrast in the basal ganglia.

## 4. Discussion

As anticipated, the extension of our previous work to include subcortical white/gray contrast in addition to cortical white/gray contrast produced both increased age-prediction accuracy and additional insight into the basis of differences in cognitive functioning. In the PING dataset, with over 900 subjects scanned on 3T scanners, the inclusion of subcortical white/gray contrast yielded an increase in the correlation of predicted and chronological age from 0.91 to 0.93 and a decrease in the mean absolute error from 613 days to 540 days. In the NIHPD dataset, with 402 subjects scanned on 1.5T scanners, the inclusion of subcortical white/gray contrast yielded a lesser increase in accuracy, with the correlation of predicted and chronological age increasing from 0.88 to 0.89, and the mean absolute error decreasing from 542 days to 500 days. Note that these results, from a single metric in a single image-type, are exceptional. For the PING data, Ball et al. (2017) produced predictions of similar, but slightly lesser, accuracy, also using a single image-type, but using multiple metrics; their correlation between predicted and chronolgical age was 0.926 and their mean absolute error was approximately 566 days. Brown et al. (2012), also using a single image-type but multiple metrics, achieved a correlation between predicted and chronological age of only 0.91, with a mean absolute error of approximately 624 days. And for the NIHPD data, Khundrakpam et al. (2015), using cortical thickness, achieved a correlation of 0.84, with a mean absolute error of 613 days. Franke et al. (2012) achieved an ostensibly superior result with the NIHPD data, but using a subsample of the scans heavily skewed toward younger subjects, *e.g.* 32% of their subjects were less than 8 years of age, whereas only 13% of ours were less then 8 years of age, and only 9% of their subjects were older than 16 years of age, whereas 15% of ours were older than 16 years of age. Comparisons across samples that differ in this way are not straightforward. Development is more rapid in younger subjects and so age prediction is more accurate; and the sparsity of the results at the upper end of the age scale in Franke et al.’s sample minimizes the negative impact of any part of the error that is due to nonlinear issues arising during adolescence. Among those studies using similar samples to those used in the current work, our methods produce more accurate predictions, including in comparison to those using multiple metrics.

And, in addition to the residuals of our age predictions reflecting individual differences in cognitive functioning, as was also the case using only cortical white/gray contrast (Lewis et al., 2018), the inclusion of the subcortical white/gray contrast measures yields a clear distinction between the relation of the age-prediction errors and VIQ versus PIQ. PIQ was shown to be positively related to the age-prediction residuals, while VIQ was shown to be negatively related to the residuals. Thus, the younger a subject’s brain appears relative to their chronological age, the higher that subject’s PIQ; and the older a subject’s brain appears relative to their chronological age, the higher that subject’s VIQ. A number of previous age-prediction studies have sought to relate their errors to cognitive function, but have generally found no relationship between the two, *e.g.* Ball et al. (2017) and Khundrakpam et al. (2015). Khundrakpam et al. (2015) report a significant relation between the age-prediction errors and the interaction of PIQ and VIQ, which is in agreement with our findings of relations in opposing directions for the relation between the age-prediction residuals and VIQ versus PIQ. But, Khundrakpam et al. (2015) found no relation between either PIQ or VIQ and their age-prediction errors. Interestingly, Erus et al. (2015) reported that individuals with brains that appeared younger than their chronological age showed inferior cognitive processing speed, and vice-versa. To the extent that processing speed correlates with PIQ, Erus et al.’s results conflict with the results here; and to the extent that processing speed correlates with VIQ, their results align with ours. But, it is unclear how processing speed relates to PIQ and VIQ; significant processing speed relations included verbal memory, but also attention, face memory, spatial memory, emotion identification, motor speed, and sensory-motor processing speed.

The inclusion of the subcortical white/gray contrast measures also provided novel insights into the brain-basis of the opposing relationships between the age-prediction residuals and PIQ versus VIQ. The top two components of a principle component analysis of the residuals of the best-fit model for the contrast data showed a categorical distinction in the brain-basis of PIQ versus VIQ, with the first component related to PIQ but not VIQ, and the second component related to VIQ but not PIQ. And the mappings of these components to the brain show that inter-individual PIQ differences involve much of the cortex but not the primary sensory areas nor any of the subcortical structures, while inter-individual VIQ differences involve the basal ganglia and the bilateral arcuate fasciculi but not the thalamus nor the rest of the cortex.

A natural interpretation of the positive relation between PIQ and the first principle component is that greater cortical contrast represents a greater degree of myelination in the fibers that interconnect wide-spread cortical regions, and that this greater myelination yields more efficient communication, and that this is a correlate of PIQ. This interpretation fits well with Jung and Haier’s 2007 Parieto-Frontal Integration Theory (P-FIT) of intelligence. The same interpretation applies to the positive relation between VIQ and the arcuate fasciculi portion of the second principle component. The arcuate fasciculi are bundles of white-matter fibers that interconnect critical components of the language network, *e.g.* Broca’s area in the inferior frontal gyrus, Wernicke’s area at the posterior of the superior temporal gyrus, and auditory cortex in the superior and middle temporal gyri. In adults, language is lateralized; in infants, language is bilateral (Perani et al., 2011; Holland et al., 2007; Broce et al., 2018; Su et al., 2018). The development of the arcuate fasciculi is reflected in cognitive ability (Schmithorst et al., 2005; Lebel and Beaulieu, 2009).

A natural interpretation of the negative relation of VIQ and the basal ganglia portion of the second principle component is similar. But, the internal architecture of subcortical structures shows substantial alterations over development, in part due to the myelination of invading fibers, in part due to maturation of fibers inter-connecting sub-nuclei, and in part due to alterations in cell structure (Takase et al., 2004; Wierenga et al., 2014; Geeraert et al., 2018). The impact of these changes on the T1 signal are not completely understood, but white/gray contrast decreases rapidly with age. Thus the negative relation between the age-prediction residuals and VIQ corresponds to a positive relation between VIQ and the state of maturation of the basal ganglia. The basal ganglia are considered to be “cognitive pattern generators” (Graybiel, 1997) regulating sequential processing, and central to sequence learning, including vocal sequences (Packard and Knowlton, 2002; Graybiel, 2005; Ölveczky et al., 2005; Dominey et al., 2003; Houk et al., 1995; Damasio, 1983; Kotz et al., 2009; Price, 2012). The accuracy of phonological processing is correlated with greater activation in the left caudate (Moro et al., 2001); the speed of phonological processing is correlated with greater activation in the left putamen (Tettamanti et al., 2005; Wildgruber et al., 2001). Syntactic processing, likewise, appears to involve the left caudate and putamen, as well as the globus pallidus (Friederici et al., 1999; Kotz et al., 2003; Friederici et al., 2003b,a; Newman et al., 2010). And, most directly related to the findings here, morphological measures of the basal ganglia have been associated with IQ (MacDonald et al., 2000; Isaacs et al., 2008; Grazioplene et al., 2015), with some specificity of the core relation being between the caudate and VIQ (Isaacs et al., 2008; Grazioplene et al., 2015).

Thus the second principle component indicates co-maturation of the arcuate fasciculi and the basal ganglia. Both the arcuate fasciculi and the basal ganglia have been linked to language, but to the best of our knowledge have not been linked to each other. That link could, of course, be a coincidence; the covariation that places the arcuate fasciculi and the basal ganglia into the same principle component might be chance. But it seems more likely that the role of the basal ganglia in phonological and syntactic processing is impacting the development of the arcuate fasciculi. Moreover, connectivity from the basal ganglia to cortical regions within the language network has been reported. In non-human primates, tracer injection studies have shown basal ganglia afferents from wide-spread regions of cortex (Yeterian and Van Hoesen, 1978; Yeterian and Pandya, 1993; Van Hoesen et al., 1981), and basal ganglia projections to a number of cortical regions in the prefrontal cortex including the supplementary motor area (SMA), the pre-SMA, and most notably to a region of the ventral premotor cortex claimed to be homologous to Broca’s area (Akkal et al., 2007; Middleton and Strick, 2000). In humans, diffusion tractography has also shown connectivity between Broca’s area and the putamen (Ford et al., 2013); tractography, however, cannot distinguish the directionality of connections, but a dynamic causal modeling analysis of functional MRI indicated unidirectional connections from the putamen to the left inferior frontal gyrus (Booth et al., 2007).

Of course, this interpretation is speculative. Speculative in part because the white/gray contrast measure is a ratio, and the intensity measures that comprise it cannot be considered in isolation; in part because the sign of a principle component is arbitrary; and in part because the co-variation that yields a principle component may stem from unrelated sources. The neurobiological properties underlying differences in the T1-weighted signal are not completely understood, but differences in the quantity and structure of myelin between white-matter and gray-matter, and the changes in myelin across development, are reflected in T1 relaxation times (Agartz et al., 1991; Barkovich et al., 1988; Barkovich, 2000; Peters, 2002). Thus the positive relation between cortical contrast and both PIQ and VIQ might stem from either relatively increased myelin within the white-matter, or relatively reduced myelin within the gray-matter, *e.g.* from a lesser degree of invasion of white-matter fibers into the cortex. Which of these is the case is unclear. Likewise, the negative relation between VIQ and contrast within the basal ganglia might stem from either relatively increased myelin within the basal ganglia, or from relatively decreased myelin in the surrounding white-matter. Again, which of these is the case is unclear. A confirmation of our interpretation of the contrast measures might be obtained from quantitative MRI, where the intensity measures in gray-matter and in white-matter are individually meaningful. But, to the best of our knowledge, no publicly available developmental data of this sort currently exists. Likewise, the conjectured impact of the development of the basal ganglia on the development of the arcuate fasciculi might be explored via analysis of longitudinal functional data in conjunction with quantitative MRI data; but, to the best of our knowledge, no publicly available developmental data of this sort currently exists. Future research should address these questions.

But regardless of the interpretation, the combination of cortical and sub-cortical white-gray contrast provides improved age-prediction accuracy over cortical white/gray contrast alone, and indicates a heretofore unseen distinction in the impact of VIQ versus PIQ on such predictions, as well as identifying the brain-basis of such a distinction.

## 5. Acknowledgments

This research has been supported by grant 316258 from the Academy of Finland (to JT), by grant ANRP-MIRI13-3388 from the Azrieli Neurodevelopmental Research Program in partnership with the Brain Canada Multi-Investigator Research Initiative (to ACE), and by grants from the Canadian Institutes of Health Research and the Natural Sciences and Engineering Research Council of Canada (to DLC). It also benefited from computational resources provided by Compute Canada (www.computecanada.ca) and Calcul Quebec (www.calculquebec.ca).

Data used in the preparation of this article were obtained from the NIH Pediatric MRI Data Repository created by the NIH MRI Study of Normal Brain Development. This is a multisite, longitudinal study of typically developing children conducted by the Brain Development Cooperative Group and supported by the National Institute of Child Health and Human Development, the National Institute on Drug Abuse, the National Institute of Mental Health, and the National Institute of Neurological Disorders and Stroke (Contract #s N01- HD02-3343, N01-MH9-0002, and N01-NS-9-2314, −2315, −2316, −2317, −2319 and −2320). A listing of the participating sites and a complete listing of the study investigators can be found at http://pediatricmri.nih.gov/nihpd/info/participating_centers.html. This manuscript reflects the views of the authors and may not reflect the opinions or views of the NIH.

Data used in the preparation of this manuscript were obtained and analyzed from the controlled access datasets distributed from the NIMH-supported Research Domain Criteria Database (RDoCdb). RDoCdb is a collaborative informatics system created by the National Institute of Mental Health to store and share data resulting from grants funded through the Research Domain Criteria (RDoC) project. This manuscript reflects the views of the authors and may not reflect the opinions or views of the NIH or of the Submitters submitting original data to RDoCdb.

## Footnotes

1 http://pingstudy.ucsd.edu/resources/neuroimaging-cores.html.

2 http://www.bic.mni.mcgill.ca/ServicesSoftware/CIVET-2-1-0-Introduction

3 We used a Matlab wrapper described by Qian et al. (2013) of Friedman’s original Fortran code.

4 The Matlab code used for constructing stratified cross-validation folds for regression is available at https://github.com/jussitohka/general_matlab

